# A single-cell transcriptomic atlas of sensory-dependent gene expression in developing mouse visual cortex

**DOI:** 10.1101/2024.06.25.600673

**Authors:** Andre M. Xavier, Qianyu Lin, Chris J. Kang, Lucas Cheadle

## Abstract

Sensory experience drives the refinement and maturation of neural circuits during postnatal brain development through molecular mechanisms that remain to be fully elucidated. One likely mechanism involves the sensory-dependent expression of genes that encode direct mediators of circuit remodeling within developing cells. However, while studies in adult systems have begun to uncover crucial roles for sensory-induced genes in modifying circuit connectivity, the gene programs induced by brain cells in response to sensory experience during development remain to be fully characterized. Here, we present a single-nucleus RNA-sequencing dataset describing the transcriptional responses of cells in mouse visual cortex to sensory deprivation or sensory stimulation during a developmental window when visual input is necessary for circuit refinement. We sequenced 118,529 individual nuclei across sixteen neuronal and non-neuronal cortical cell types isolated from control, sensory deprived, and sensory stimulated mice, identifying 1,268 unique sensory-induced genes within the developing brain. To demonstrate the utility of this resource, we compared the architecture and ontology of sensory-induced gene programs between cell types, annotated transcriptional induction and repression events based upon RNA velocity, and discovered Neurexin and Neuregulin signaling networks that underlie cell-cell interactions via *CellChat*. We find that excitatory neurons, especially layer 2/3 pyramidal neurons, are highly sensitive to sensory stimulation, and that the sensory-induced genes in these cells are poised to strengthen synapse-to-nucleus crosstalk by heightening protein serine/threonine kinase activity. Altogether, we expect this dataset to significantly broaden our understanding of the molecular mechanisms through which sensory experience shapes neural circuit wiring in the developing brain.

## Introduction

The precise connectivity of neural circuits in the mammalian brain arises from a convergence of genetic and environmental factors spanning embryonic and postnatal stages of development. Brain circuits are first assembled *in utero* via the formation of an overabundance of nascent synaptic connections between potential neuronal partners, then later remodeled, or *refined*, postnatally through the strengthening of some of these synapses and the elimination of others^1,2^. The selective retention and maturation of a subset of initially formed synapses equips the brain with an interconnected network of circuits optimized to facilitate neurological function and plasticity across the lifespan. Furthermore, impairments in postnatal phases of synaptic remodeling and refinement are increasingly appreciated to contribute to a host of neurodevelopmental conditions such as autism and schizophrenia^3–5^. Thus, elucidating the mechanisms underlying circuit refinement in the early postnatal brain is important from both basic and translational perspectives.

While much emphasis has been placed on defining the intrinsic genetic mechanisms that govern embryonic stages of brain development, such as neurogenesis, cell migration, and synapse formation, less is known about how environmental triggers that come online after birth shape the maturation of neural circuits as they emerge. A prime example of environmental cues impacting circuit development can be seen in the role of sensory experience in driving the refinement of neural circuitry within the visual system of the mouse, a process that takes place around the third week of life^6^. Specifically, between postnatal days (P)20 and P30, visual experience promotes the structural and functional refinement and maturation of synaptic connections between excitatory thalamocortical neurons within the dorsal lateral geniculate nucleus (dLGN) of the thalamus and their postsynaptic targets in layer 4 of primary visual cortex (V1)^7–10^. Importantly, blocking visual experience during this developmental window by rearing mice in complete darkness significantly impedes the maturation and eventual function of visual circuits, whereas blockade of experience outside of this time frame does not have a strong observable effect on circuit wiring^11–13^. Thus, sensory experience drives the developmental refinement of thalamocortical circuits selectively during a defined window of postnatal brain development in the mouse visual cortex.

While the visual system has provided numerous essential insights into functional aspects of synaptic refinement and plasticity, our understanding of the molecular mechanisms that mediate the influence of visual experience on circuit wiring remains limited. One likely mechanism linking experience to circuit development in V1 is the induction of gene programs in neurons in response to sensory-driven neuronal activity, a process termed *activity-dependent transcription*^14^. In this process, synaptic innervation onto a given neuron drives the influx of Ca^2+^ into the cell through ionotropic glutamate receptors and L-type Ca^2+^ channels^15^. This influx of Ca^2+^ initiates intracellular signaling cascades that phosphorylate, and thereby activate, transcription factors in the nucleus, such as CREB and MEF2^16,17^. Within the first hour of neuronal activation, these factors induce the expression of immediate early genes (IEGs), many of which encode a separate set of transcription factors, including the well-established IEG *Fos*^18^. During a second wave of activity-dependent transcription which typically occurs between two and six hours following neuronal activation, IEGs bind a subset of genomic promoters and enhancers to drive the expression of a separate cohort of genes (late-response genes, LRGs) encoding direct mediators of synaptic remodeling, such as the secreted neurotrophin *Bdnf*^19,20^. This two-wave pattern of activity-dependent transcription encompassing the early expression of transcriptional regulators (i.e. IEGs) followed by the later expression of molecules that act locally at individual synapses (i.e. LRGs) is likely to contribute to the refinement of visual circuits at the molecular level. Although a detailed analysis of sensory-driven transcription during the critical window of circuit development in V1 has not yet been established, work in adult animals suggests that the gene programs induced by experience in this brain region are highly cell-type-specific, reflecting the capacity of activity-dependent genes to shape cellular function in a precise manner and necessitating investigation at cell-type resolution^21–23^. Thus, genes that are induced by neurons in response to sensory experience are promising candidates to mediate circuit development during a period of sensory-dependent refinement in V1.

Given the potential of sensory-induced gene programs to represent a key molecular link between visual experience and circuit development, we reasoned that a comprehensive atlas of sensory-dependent gene expression in developing V1 would provide useful insights into how circuits mature in response to environmental cues at the molecular level. To this end, we leveraged a sensory deprivation and stimulation paradigm, the late-dark-rearing (LDR) paradigm, to dampen or increase visual experience between P20 and P27, when sensory-dependent synapse remodeling begins and peaks, respectively. We then performed single-nucleus RNA-sequencing (snRNAseq) on V1 tissue across six conditions encompassing normally reared, sensory deprived, and four cohorts of sensory stimulated mice, followed by differential gene expression analysis to identify transcripts that exhibited significantly greater RNA abundance in cells from stimulated versus unstimulated animals. We then further demonstrated the utility of the dataset by comparing sensory-dependent gene programs between cell types, characterizing transcriptional induction and repression events based upon RNA velocity, and predicting molecular interactions between cell types using the computational tool *CellChat*. We expect this dataset, which describes sensory-driven changes in gene expression across 118,529 nuclei representing sixteen distinct types of brain cells, to be a valuable resource for investigators interested in uncovering the molecular basis of sensory-dependent synapse remodeling and plasticity in the developing brain.

## Results

### A visual deprivation and stimulation paradigm for capturing sensory-induced transcripts

To characterize the gene programs that are elicited by sensory experience during a critical period of postnatal brain development, we harnessed a dark-rearing method to manipulate visual experience in mice in a temporally restricted manner. In this paradigm, mice were initially reared according to a standard 12-hour light/12-hour dark cycle (normal rearing, NR), the environment maintained in most animal facilities, before being placed in complete darkness at P20, the beginning of sensory-dependent visual circuit refinement. Mice were then maintained in a completely dark environment for 24 hours a day across a seven-day period. At P27, when sensory-dependent refinement peaks, one cohort of dark-reared mice was sacrificed in the dark without re-exposure to light, and V1 tissue was collected. Other cohorts of dark-reared mice were acutely re-exposed to white light at P27 for varying amounts of time following the week-long period of darkness, a manipulation that leads to the acute and robust activation of circuit refinement and plasticity in both the dLGN and V1^6,11,24^. Thus, this late-dark-rearing (LDR) paradigm allowed us to assess the impact of (1) sensory deprivation, and (2) sensory stimulation on gene expression selectively during a time when such experience is necessary for neural circuit refinement (Fig. 1A).

**Figure 1.**
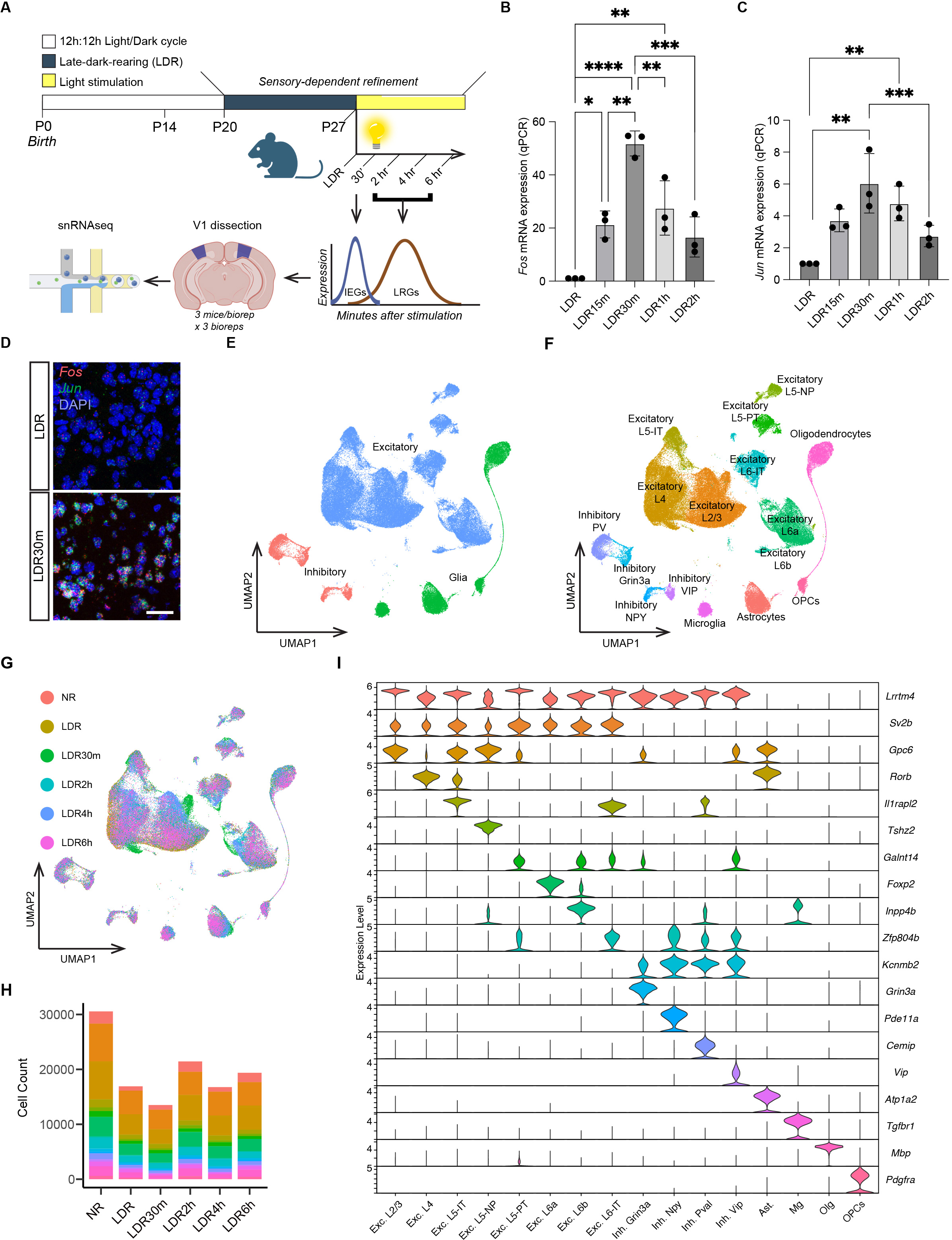
Experimental design and introduction to the single-nucleus RNA-sequencing dataset. (A) Schematic describing the late-dark-rearing (LDR) paradigm and the workflow of the single-nucleus RNA-sequencing (snRNAseq) experiments. (B) Quantification of *Fos* mRNA expression in sensory deprived (LDR) mice and in mice acutely exposed to light for between 15 minutes and 2 hours, with stimulation timepoints labeled as follows: LDR15m (15 min of light), LDR30m (30 min), LDR1h (1 hour), and LDR2h (2 hours). *Fos* expression assessed by qPCR and normalized to *Gapdh* expression. Values plotted are additionally normalized to the LDR condition. (C) qPCR quantification of *Jun* mRNA expression (normalized to *Gapdh*) in V1 across all timepoints. Data obtained by qPCR and values plotted are normalized to LDR. (D) Example confocal images of V1 in sections from a sensory deprived mouse (LDR) and a mouse re-exposed to light for thirty minutes (LDR30m). *Fos* mRNA (red), *Jun* mRNA (green), and DAPI (blue). Scale bar, 44 μm. (E) UMAP plot illustrating the 118,529 nuclei in the dataset categorized by general cell class: excitatory neurons (periwinkle), inhibitory neurons (salmon), and glia (green). (F) UMAP plot with all 16 clusters colored and labeled by cell type. (G) UMAP plot with cells colored by condition according to the legend on the left. Note that cluster composition is largely unaffected by sensory deprivation or stimulation. (H) Numbers of cells of each type included in the final dataset across all conditions. See also Table 1. (I) Violin plot demonstrating the enrichment of markers used to assign nuclei in the dataset to distinct cell types. Top enriched gene per cluster given on the Y-axis on the right, normalized FPKM expression given on the Y-axis on the left, and cluster identity shown on the X-axis. For (B) and (C), n = 3 mice per condition; One-way ANOVA followed by Tukey’s post hoc test: *p < 0.05; **p < 0.01; ***p < 0.001; ****p < 0.0001.

Previous studies of sensory-dependent transcription in the visual system have focused on analyzing gene expression at two timepoints following the re-exposure of adult mice to light after dark-rearing (compared to unstimulated, sensory deprived mice): one hour, when immediate-early genes (IEGs) have been proposed to peak, and four hours, when late-response genes (LRGs) are highly induced^21^. However, given that these studies were performed in adult mice, we first sought to confirm that these time points were also optimal for capturing sensory-dependent gene expression during postnatal development. To do so, we performed qPCR and single-molecule fluorescence *in situ* hybridization (smFISH) in parallel on V1 tissue after subjecting mice to the LDR paradigm described above, and we assessed the expression of the canonical IEGs *Fos* and *Jun* as a read-out for the timing of sensory-dependent transcription. We found that the expression of both IEGs was increased as early as fifteen minutes after light re-exposure (i.e. acute sensory stimulation), and that this increase in *Fos* and *Jun* persisted for at least two hours after stimulation. Within this time frame, the peak of *Fos* and *Jun* expression occurred not at one hour but at thirty minutes after light re-exposure (Fig. 1B-D). Thus, we included a thirty-minute stimulation timepoint to capture IEGs in our experiments. We also included three additional stimulation timepoints at which we expected to capture the bulk of LRGs: two hours, four hours, and six hours. An additional benefit of including three late-response conditions is that it allowed us to derive insights into the dynamics of sensory-dependent gene programs on a broader scale. Altogether, our finalized dataset includes cells from mice according to the following six conditions: normally reared (NR) mice at P27, mice reared in complete darkness from P20 to P27 (LDR), and mice reared in complete darkness between P20 and P27 then acutely re-exposed to light for 30 minutes (LDR30m), two hours (LDR2h), four hours (LDR4h), or six hours (LDR6h). To our knowledge, this is the most extensive time course of sensory-dependent gene expression at single-cell resolution generated to date.

### Mapping sensory-dependent gene expression in the developing cortex

To map sensory-dependent changes in gene expression across cortical cell types in an unbiased manner, we performed single-nucleus RNA-sequencing (snRNAseq; 10X Genomics) on V1 tissue bilaterally micro-dissected from mice following the LDR paradigm described above (Fig. 1A). We sequenced individual nuclei rather than whole cells based upon our interest in capturing nascent transcriptional events that are acutely induced by experience. Three biological replicates were performed for each condition, with each replicate being made up of cells pooled from the visual cortices of three animals to increase yield. Biological replicates were collected, isolated, and processed independently on different days to control for batch effects. After next-generation sequencing, the data were mapped to the mouse genome and quality control was performed to remove putative doublets, unhealthy or dying cells, and droplets containing ambient RNA from the dataset using Seurat, DoubletFinder, and DecontX packages in R^25–27^. Data were then integrated across biological replicates and conditions for downstream analysis within Seurat. The final dataset includes 118,529 nuclei across 16 distinct cell clusters representing eight excitatory neuron subtypes, four inhibitory neuron subtypes, and four glial subtypes (Fig. 1E-G). The excitatory populations captured include layer 4 (L4) pyramidal (PYR) neurons, layer 2/3 (L2/3) PYR neurons, three populations of layer 5 (L5) neurons, and three populations of layer 6 (L6) neurons. Inhibitory populations sequenced include Grin3a-enriched neurons (some of which also express SST markers), parvalbumin (PV) neurons, VIP neurons, and neurons expressing Npy, which include neurogliaform cells (also positive for Lamp5) and a subset of SST neurons. Glial populations sequenced include astrocytes, oligodendrocytes, oligodendrocyte precursor cells (OPCs), and microglia (Fig. 1H,I). Cell type assignments were based upon the presence of marker genes identified previously^21,28^. The numbers of cells within each cell class included in the dataset are given in Table 1.

**Table 1.** Number of each cell type represented in the dataset. displays the number of cells included in the final dataset by condition and cell type. Cell types listed in alphabetical order from top to bottom.

### Sensory deprivation upregulates a cohort of genes in excitatory neurons

With this dataset in hand, we set out to understand how manipulating sensory experience impacts the transcriptional states of cells in V1 during development. To this end, we utilized the DEseq2 function within Seurat to identify transcripts that were significantly differentially expressed (differentially expressed genes, DEGs; false discovery rate (FDR) < 0.05) between each condition for each cell type, beginning with a comparison of gene expression in normally reared (NR) mice at P27 versus sensory deprived (i.e. LDR) mice at the same age. This analysis revealed changes in gene expression meeting a minimum threshold of log_2_(1.5) fold change in excitatory neurons following dark-rearing compared to NR mice. Specifically, when the DEG analysis was applied to all excitatory neuron clusters in aggregate, 52 genes (e.g. the transcription factor *Stat4* and the cytoskeletal regulator *Clmn*) were found to be less highly expressed in the NR condition compared to LDR mice, indicating that depriving mice of light *increased* the expression of a define cohort of genes (Fig. 2A). Among excitatory neuron clusters, the subtypes that exhibited the largest numbers of gene expression changes following LDR were L2/3 PYR neurons (86 genes upregulated after dark-rearing and 7 genes downregulated; Fig. 2B) followed by neurons in the L6a cluster (21 genes upregulated after dark-rearing; Fig. 2C). In L2/3 neurons, genes more highly expressed in the LDR condition included factors such as *Tspan11* and *Gpc3* which are involved in cellular dynamics and migration^29,30^. Interestingly, the seven genes that were more highly expressed in the NR condition included known activity-regulated factors such as the neurotrophin *Bdnf* and the nuclear orphan receptor *Nr4a1*. In contrast to excitatory neurons, only one gene, the synaptically localized long non-coding RNA *Gm45323*, was less highly expressed in inhibitory neurons in the NR compared to the LDR condition^31^ (Fig. 2D). These findings suggest that excitatory neurons are more sensitive to the effects of sensory deprivation than inhibitory neurons at the transcriptional level, and indicate that sensory deprivation tends to increase, rather than decrease, gene expression in excitatory neurons of V1.

**Figure 2.**
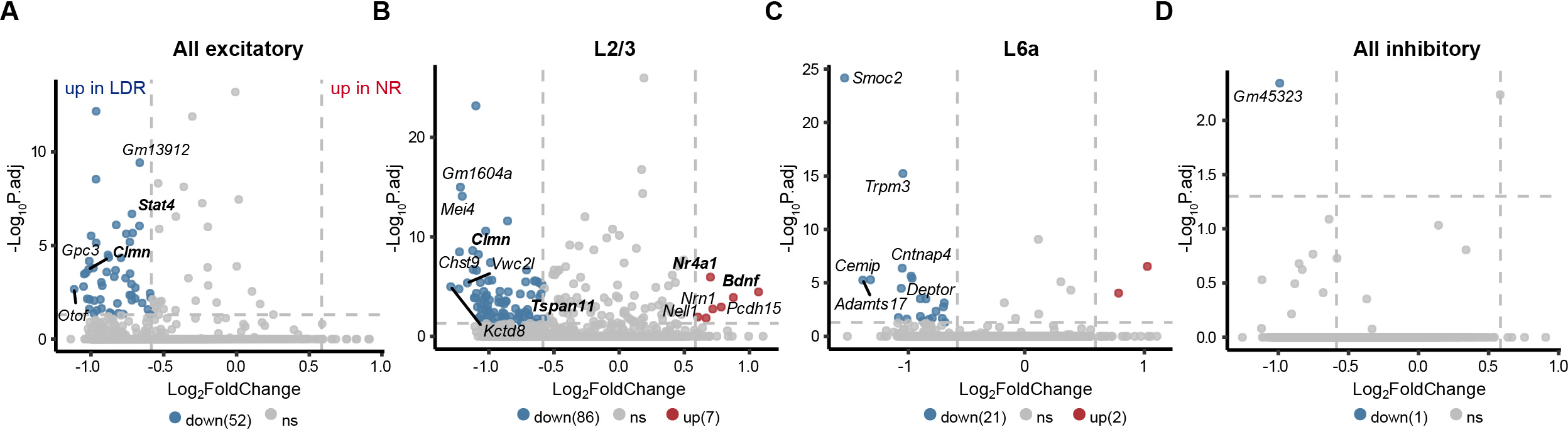
Sensory deprivation upregulates a cohort of genes in excitatory neurons. (A) Volcano plot demonstrating transcripts that were significantly differentially expressed (differentially expressed genes, DEGs) in aggregated excitatory neuron clusters after LDR compared to normally reared (NR) control mice. Y-axis, negative Log(10) adjusted p value (threshold of p.adj < 0.05 indicated by dashed horizontal line). X-axis, Log(2) fold change (threshold of log_2_(1.5) indicated by dashed vertical lines). Red, genes that are more highly expressed in the NR condition (up in NR). Blue, genes that are more highly expressed in the LDR condition (up in LDR). (B) Volcano plot of DEGs altered by sensory deprivation in excitatory L2/3 neurons. (C) Volcano plot of DEGs altered by sensory deprivation in L6a neurons. (D) Volcano plot of DEGs altered by sensory deprivation in aggregated inhibitory clusters.

### Excitatory and inhibitory neurons mount shared and distinct transcriptional responses to sensory stimulation

We next compared DEGs between each light re-exposure timepoint (LDR30m, LDR2h, LDR4h, and LDR6h) and the sensory deprived LDR condition for all 16 cell clusters in isolation, and for inhibitory and excitatory populations after aggregation. These experiments revealed bidirectional changes in gene expression at every timepoint analyzed within most cell types, yielding a total number of 1,268 unique genes that are upregulated at any stimulation timepoint compared to the LDR condition (Table 2). These genes included numerous previously identified activity-dependent IEGs, such as the AP1 factors *Fos* and *Jun*, the neuron-specific IEG *Npas4*, and the *Nr4a* and *Egr* families of TFs that are induced by various extracellular stimuli including synaptic innervation. Interestingly, although AP1 transcription factors are broadly considered to be IEGs, *Jund* and *Junb* both exhibited a pattern of induction more consistent with an LRG identity, peaking at LDR2h rather than LDR30m (Fig. 3A). Among all cell types analyzed, sensory experience elicited the most robust gene expression changes in L2/3 and L4 excitatory neurons, a result that we validated by performing smFISH for the IEGs *Fos* and *Nr4a1* in these layers (Fig. 3B-E). Both of these excitatory populations are innervated by projection neurons outside of V1, with L2/3 neurons receiving top-down information from other cortical areas and L4 neurons receiving bottom-up information from the visual thalamus that is strongly driven by sensory experience. That L2/3 PYR neurons exhibit the largest number of transcriptional changes as a result of light re-exposure is consistent with a recent report identifying L2/3 cells as being particularly sensitive to sensory experience during postnatal development^32^. In addition to IEGs, we also identified cohorts of genes that were preferentially upregulated at LDR2h, LDR4h, or LDR6h, fitting the expected profile of LRGs (Fig. 3F). Assessing changes in gene expression following light re-exposure in aggregated excitatory and inhibitory neuron populations revealed that the most robust change in gene expression for excitatory neurons (233 genes upregulated) occurred at the 30-minute timepoint, while the most robust change in inhibitory neurons (82 genes upregulated) occurred six hours after light re-exposure (Fig. 3G). This observation could reflect a temporal trajectory in which excitatory neurons are more strongly impacted by sensory stimulation first, with inhibitory neurons responding later.

**Figure 3.**
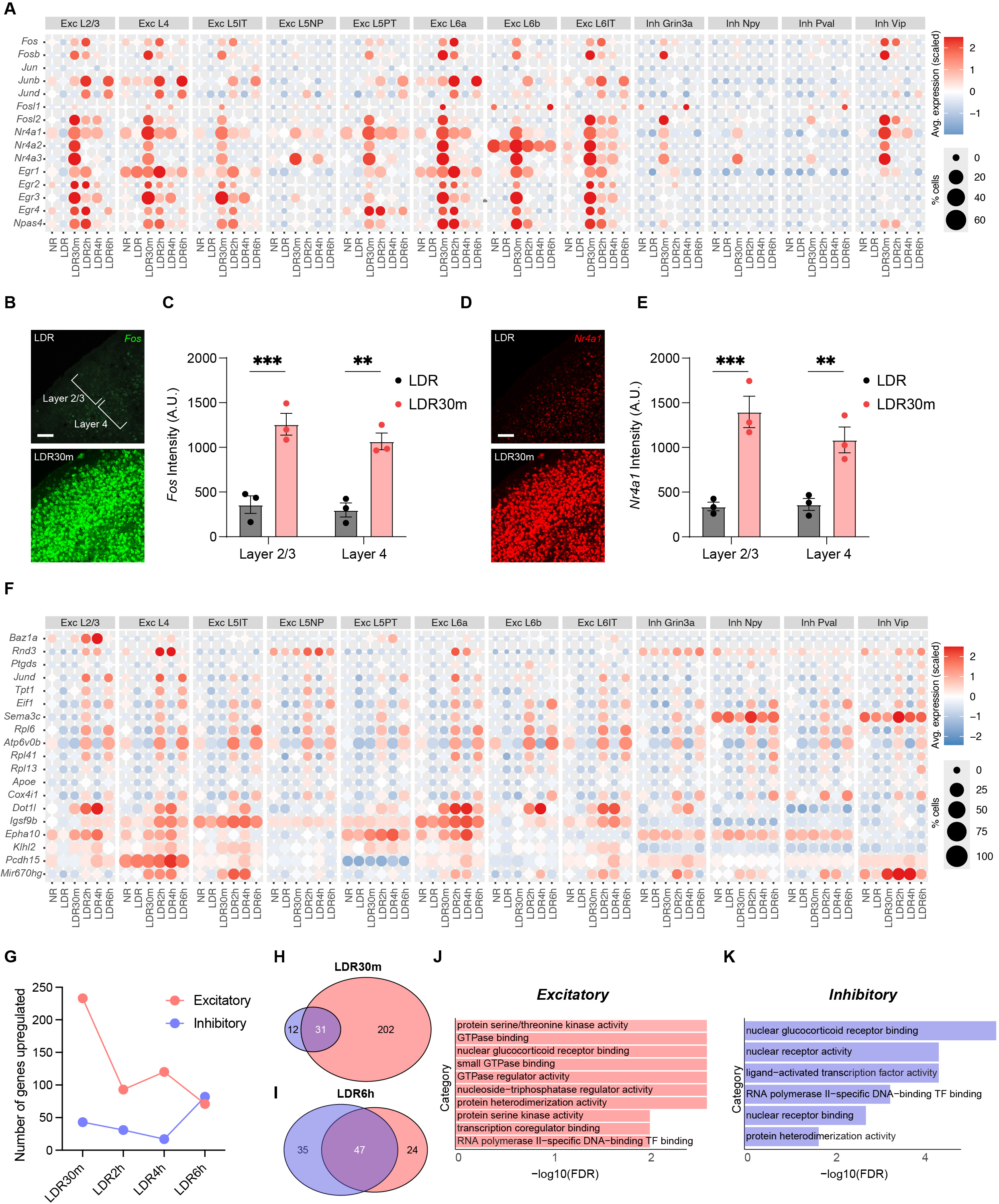
Excitatory and inhibitory neurons mount shared and distinct responses to sensory stimulation. (A) Bubble plot illustrating the induction of canonical immediate-early genes (IEGs) across timepoints and cell types. Color indicates relative expression level according to the scale on the right. Size of circle represents the percentage of cells expressing the gene. (B) Confocal images of V1 in late-dark-reared (LDR) mice and in mice re-exposed to light for 30 min (LDR30m) subjected to single molecule fluorescence *in situ* hybridization (smFISH) to label *Fos* mRNA. Scale bar, 100 μm. (C) Quantification of *Fos* expression (arbitrary units, A.U.) in L2/3 and L4 of V1 in LDR and LDR30m mice. Two-Way ANOVA with Tukey’s post hoc test. **p<0.01, ***p<0.001; n = 3 mice/condition. (D) Confocal images of V1 in LDR and LDR30m mice subjected to smFISH to label *Nr4a1* mRNA. Scale bar, 100 μm. (E) Quantification of *Nr4a1* expression in L2/3 and L4 in LDR and LDR30m mice. Two-Way ANOVA with Tukey’s post hoc test. **p<0.01, ***p<0.001; n = 3 mice/condition. (F) Bubble plot demonstrating late-response gene (LRG) expression across cell types and conditions. Scaled expression indicated on the right. (G) Graph displaying the numbers of genes significantly upregulated at each stimulation timepoint (compared to LDR control) across conditions for aggregated excitatory (salmon) and inhibitory (periwinkle) neurons. (H) Venn diagram demonstrating overlap between sensory-dependent gene programs in excitatory (salmon) versus inhibitory (periwinkle) neurons at LDR30m. (I) Venn diagram demonstrating overlap between sensory-dependent gene programs in inhibitory versus excitatory neurons at LDR6h. (J) Gene ontology (GO) categories enriched among genes upregulated in excitatory neurons at LDR30m. (K) GO categories enriched among genes upregulated in inhibitory neurons at LDR30m.

**Table 2.** Differentially expressed genes identified in each cell type. includes all significantly differentially expressed genes identified by DEseq2 across cell types and conditions. Cell types and conditions listed in alphabetical order from top to bottom.

Studies in adult mice have suggested that IEGs tend to be conserved between cell types while LRGs are more likely to be cell-type-specific. Thus, we next assessed the overlap between the DEGs that were upregulated by stimulation at each time point across aggregated inhibitory and excitatory clusters. Unexpectedly, of the 233 genes that are upregulated in excitatory neurons following light re-exposure at LDR30m, only 31 (13.3%) were also upregulated in inhibitory cells at the same timepoint. Conversely, 47 (66%) of the 71 genes upregulated in excitatory neurons at LDR6h were shared with inhibitory neurons (Fig. 3H,I). These findings suggest one or both of two possibilities: (1) the LRG programs within these cell types have more in common than earlier waves of sensory-induced transcription; and/or (2) inhibitory neurons may respond more slowly to visual stimulation than excitatory neurons. The latter possibility is consistent with the finding that excitatory neurons are more sensitive to sensory deprivation than inhibitory neurons at the transcriptomic level (Fig. 2).

The commonalities and distinctions between the gene programs induced by experience in excitatory versus inhibitory neurons were reflected in the functional classifications of the sensory-induced genes identified in each cell type. For example, at LDR30m, DEGs in both classes were enriched for gene ontology (GO) categories such as *RNA polymerase II-specific DNA-binding transcription factor binding*, reflecting the sensory-induced expression of members of the Nr4a family of nuclear orphan receptors in both excitatory and inhibitory cells. Conversely, GO categories related to *GTPase binding* and *GTPase regulator activity*, including the Rho GTPase guanine nucleotide exchange factors (RhoGEFs) *Arhgef3* and *Plekhg5*, were selectively upregulated in excitatory neurons at this timepoint (Fig. 3J,K). This observation suggests that excitatory neurons may undergo structural remodeling as a result of sensory-dependent transcription in a manner that is unique from inhibitory neurons, although these transcriptional events would likely take longer than six hours to elicit a functional effect. Overall, these data indicate that the gene programs induced in excitatory and inhibitory neurons downstream of sensory stimulation exhibit partial overlap at each time point analyzed, with the amount and nature of overlap varying significantly by condition. These observations support the utility of the snRNAseq dataset for uncovering transcripts that are upregulated by sensory experience across multiple cell types as well as the transcripts that are induced by experience in a cell-type-specific manner.

### Comparison of sensory-induced genes in L2/3 and L4 excitatory neurons reveals a shared protein kinase signature and divergent axon guidance pathways

Given that L2/3 and L4 neurons were among the most strongly impacted by sensory experience, we next compared sensory-driven gene programs between these excitatory subpopulations. We first asked whether sensory-dependent gene programs in L2/3 and L4 neurons share a general architecture by quantifying the numbers of genes that were upregulated in both cell types following visual stimulation at each experimental timepoint. For both cell classes, LDR30m was the time point at which the highest numbers of genes (303 genes in L2/3 neurons and 239 genes in L4 neurons) were upregulated, followed by LDR4h, when 210 and 124 genes were upregulated in L2/3 and L4 neurons, respectively (Fig. 4A-F). This observation suggested that, among the three later timepoints analyzed, LDR4h was likely the peak of late-response, sensory-dependent transcription in L2/3 and L4 neurons. We next assessed the overlap between the gene programs induced by L2/3 and L4 neurons at each timepoint. As expected, these gene programs exhibited substantial overlap. For example, of the combined 349 genes upregulated at LDR30m across both cell classes, 193 (or 55%) were induced in both cell types (Fig. 4G). Varying degrees of overlap were also observed between sensory-dependent gene programs at the later timepoints as follows: 31% overlap at LDR2h, 36% at LDR4h, and 52% at LDR6h (Fig. 4H-J). Thus, L2/3 and L4 neurons mounted both shared and distinct responses to sensory experience that were most robust at LDR30m followed by LDR4h.

**Figure 4.**
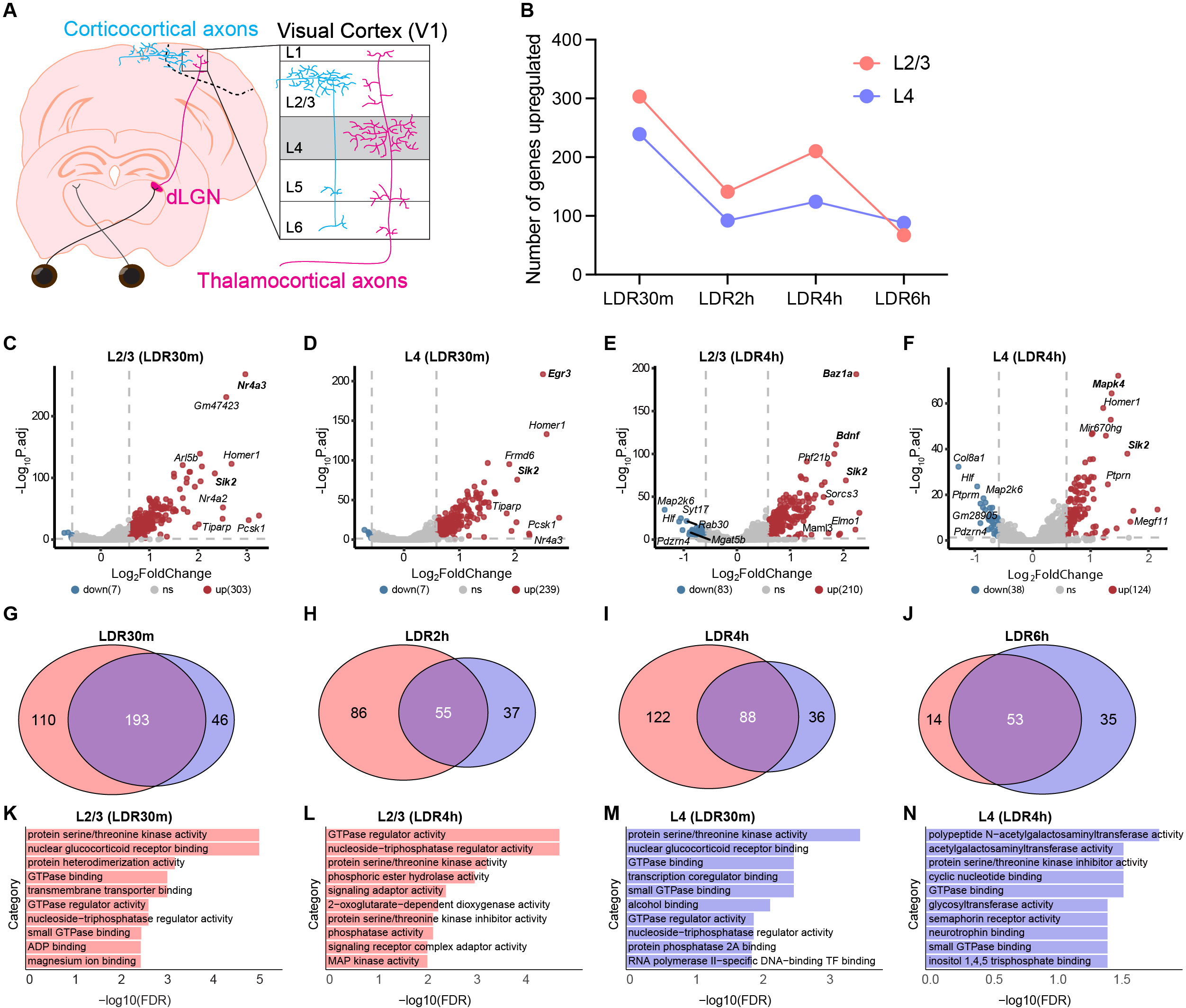
Comparison of sensory-driven gene expression in L2/3 and L4 excitatory neurons reveals a shared protein kinase signature and divergent axon guidance pathways. (A) Schematic of the pathway from the retina to primary visual cortex (V1) in the mouse. L2/3 neurons principally receive ‘top-down’ input from other regions of cortex (blue), while L4 neurons receive “bottom-up” inputs from visual thalamus (magenta). (B) Graph displaying the numbers of genes significantly upregulated at each stimulation timepoint (compared to late-dark-reared [LDR] control) across conditions for L2/3 (salmon) and L4 (periwinkle) neurons. (C) Volcano plot illustrating genes that were significantly upregulated (red) or downregulated (blue) in L2/3 neurons after 30 minutes of light re-exposure following LDR. (D) Volcano plot illustrating genes that were significantly upregulated (red) or downregulated (blue) in L4 neurons after 30 minutes of light re-exposure following LDR. (E) Volcano plot illustrating genes that were significantly upregulated (red) or downregulated (blue) in L2/3 neurons after 4 hours of light re-exposure following LDR. (F) Volcano plot illustrating genes that were significantly upregulated (red) or downregulated (blue) in L4 neurons after 4 hours of light re-exposure following LDR. (G) Venn diagram displaying overlap between upregulated genes identified in L2/3 (salmon) versus L4 neurons (periwinkle) at the LDR30m timepoint. (H) Venn diagram displaying overlap between upregulated genes in L2/3 versus L4 neurons at the LDR2h timepoint. (I) Venn diagram displaying overlap between upregulated genes identified in L2/3 versus L4 neurons at the LDR4h timepoint. (J) Venn diagram displaying overlap between upregulated genes identified in L2/3 versus L4 neurons at the LDR6h timepoint. (K) Gene ontology (GO) analysis of genes upregulated by light in L2/3 neurons at LDR30m. (L) GO analysis of genes upregulated by light in L2/3 neurons at LDR4h. (M) GO analysis of genes upregulated by light in L4 neurons at LDR30m. (N) GO analysis of genes upregulated by light in L4 neurons at LDR4h.

We next investigated the nature of the sensory-dependent gene programs induced by both cell types by performing GO analysis on the sets of genes that were upregulated at LDR30m or LDR4h. As expected based upon the overlap in the gene programs induced in L2/3 and L4 neurons (Fig. 4G-J), several of the same GO categories emerged for both cell types. These shared functional classifications included *GTPase binding* (likely reflecting mechanisms of cytoskeletal remodeling) and *nuclear receptor binding* (associated with activity-dependent transcription factors), but the *protein serine/threonine kinase activity* category was particularly prominently represented. In L2/3 neurons, this category was the most highly enriched functional classification of genes revealed by GO analysis at LDR30m and the third most enriched at LDR4h (Fig. 4K,L). Similarly, *protein serine/threonine kinase activity* is also the most enriched classification of upregulated genes in L4 neurons at LDR30m (although not at LDR4h; Fig. M,N). Thus, genes encoding protein serine/threonine kinases were among the most strongly induced genes following sensory stimulation in both L2/3 and L4 neurons, and they were induced as early as LDR30m suggesting that their expression adheres to an IEG-like pattern. On the contrary, these kinases were largely not induced by light re-exposure in inhibitory neurons at LDR30m or LDR4h (Fig. 3K), suggesting that they may be particularly important for mediating sensory-dependent plasticity in excitatory cells.

While genes within the enriched *protein serine/threonine kinase activity* category included those that encode intracellular molecules that regulate a wide range of cellular processes, in neurons, these pathways are specialized to convey information about changes at the cell membrane (and at synapses in particular) to the nucleus to shape gene expression following synaptic innervation^33,34^. For example, the Extracellular signal-regulated (ERK)-family kinases *Mapk4* and *Mapk6* were strongly upregulated by sensory stimulation at LDR30m in both L2/3 and L4 neurons, with *Mapk4* representing one of the most highly induced genes in L4 neurons at LDR4h. Likewise, the related Salt-inducible kinases, *Sik1-3*, were among the most highly upregulated genes in both cell types at all timepoints analyzed (Fig. 4C-F). Notably, intracellular signaling molecules, including ERK and Sik family kinases, interact with numerous IEG transcription factors identified in the dataset^35–38^. Thus, genes upregulated by sensory experience in L2/3 and L4 neurons share a protein kinase signature that we predict may strengthen synapse-nucleus crosstalk following sensory stimulation principally in excitatory neurons.

We next interrogated differences between the sensory-dependent gene programs in L2/3 and L4 neurons by performing GO analysis on the gene sets that were uniquely induced in each cell type. An interesting pattern to emerge was the differential induction of two axon guidance pathways within these populations: the ephrin pathway (including ephrin receptors *Ephb3* and *Epha10*) in L2/3 neurons and the semaphorin pathway (including the semaphorin co-receptors *Plxna4* and *Nrp1*) in L4 neurons (Table 2). Both of these pathways mediate the migration of neuronal axons and the establishment of synapses within target zones based upon ephrin and semaphorin ligand expression, and have been implicated in establishing retinotopy in the developing visual system^39–42^. These findings suggest that sensory experience may elicit axonal remodeling and/or presynaptic plasticity by inducing the expression of members of two distinct signaling families, ephrins and semaphorins, in L2/3 and L4 neurons, respectively. This result is consistent with the different projection patterns of these two cell types.

### Sensory-dependent gene induction and repression dynamics in L2/3 and L4 neurons revealed by RNA velocity

Increases in RNA abundance following sensory stimulation are often interpreted to reflect the new transcription of genes. However, RNA abundance can be influenced by many mechanisms beyond transcription, such as changes in the stability or degradation of the mRNA. As single-cell transcriptomic approaches have evolved over the past decade, new computational methods for disentangling transcriptional induction events from other modes of gene regulation have become available. We applied one such principle, RNA Velocity, to explicitly characterize genes whose expression is likely to be upregulated by sensory stimulation via a transcriptional mechanism. Briefly, this approach estimates transient cell state transcriptional dynamics based upon the relative abundant of nascent (unspliced) and mature (spliced) mRNAs.

Given that the earliest versions of single-cell velocity (scVelo) analysis modules exhibited limited performance in identifying multiple rate kinetics (MURK) genes^43,44^, which can exhibit rapid and complex changes at the transcriptional level, in our dataset, we applied a newer approach, UniTVelo, to assess RNA velocity in L2/3 and L4 neurons across LDR and light re-exposure timepoints^45^. This framework utilizes a radial basis function (RBF) model to overcome the limitations caused by the linear assumptions employed by scVelo. UniTVelo’s advanced approach allows for a more nuanced understanding of gene expression patterns, accurately capturing the dynamics of genes that display variable transcription rates at different stages of a cellular process.

Using UniTVelo, we first analyzed the architecture of transcript maturation for each predicted cell state transition: LDR to LDR30m, LDR30m to LDR2h, LDR2h to LDR4h, and LDR4h to LDR6h. These comparisons revealed strong signatures of both transcriptional induction and repression in L2/3 and L4 neurons with a stereotyped pattern shared by both cell types (Fig. 5A). For example, between LDR and LDR30m, relatively large numbers of genes in each cell type (204 and 161 genes in L2/3 and L4 cells, respectively) exhibited transcriptional induction with only very few genes exhibiting repression at this timepoint. On the contrary, between LDR30m and LDR2h, the majority of significantly altered genes were repressed, not induced. Between LDR2h and LDR4h, most altered genes were induced although many genes were also repressed. Finally, between LDR4h and LDR6h, the majority of altered genes in each cell type exhibited a repressed profile (Fig. 5B,C). These results are in line with the canonical view of stimulus-dependent gene programs involving primarily two waves of transcription: an IEG wave peaking at LDR30m and a LRG wave peaking at LDR4h.

**Figure 5.**
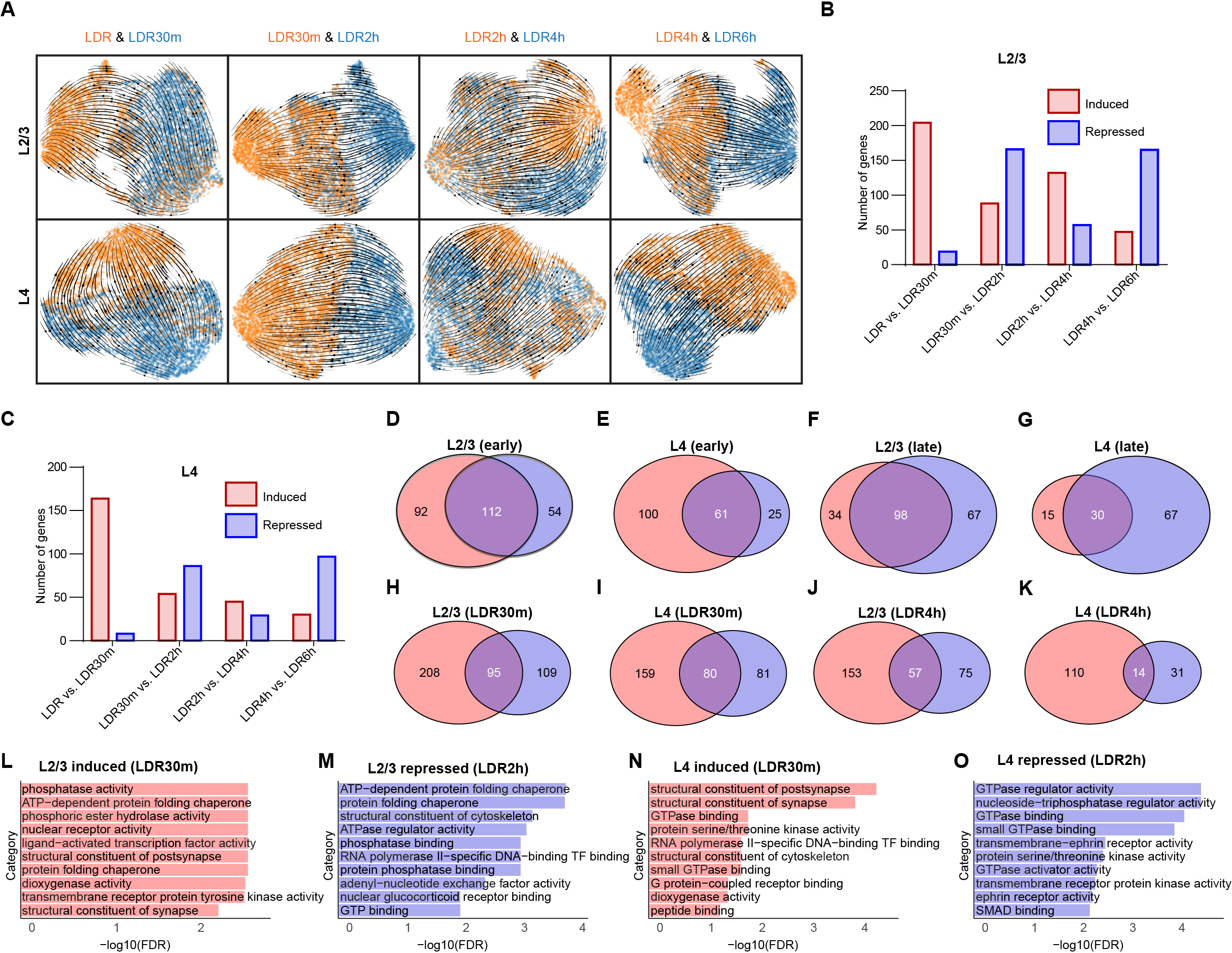
Transcriptional induction and repression events in L2/3 and L4 neurons revealed by RNA Velocity. (A) UMAP plots generated based upon RNA Velocity displaying transcriptional dynamics across each cell-state transition. L2/3 neurons, top row, L4 neurons, bottom row. (B) Bar graph displaying the total numbers of induced (red) and repressed (blue) genes across each cell-state transition in L2/3 neurons. (C) Bar graph displaying the total numbers of induced (red) and repressed (blue) genes across each cell-state transition in L4 neurons. (D) Venn diagram displaying overlap between the genes induced at LDR30m (red) and the genes that are repressed between LDR30m and LDR2h (blue) in L2/3 neurons. (E) Venn diagram displaying overlap between the genes induced at LDR30m (red) and the genes that are repressed between LDR30m and LDR2h (blue) in L4 neurons. (F) Venn diagram displaying overlap between the genes induced between LDR2h and LDR4h (red) and the genes that are repressed between LDR4h and LDR6h (blue) in L2/3 neurons. (G) Venn diagram displaying overlap between the genes induced between LDR2h and LDR4h (red) and the genes that are repressed between LDR4h and LDR6h (blue) in L4 neurons. (H) Overlap between upregulated DEGs and induced genes in L2/3 neurons at LDR30m. (I) Overlap between upregulated DEGs and induced genes in L4 neurons at LDR30m. (J) Overlap between upregulated DEGs and induced genes in L2/3 neurons at LDR4h. (K) Overlap between upregulated DEGs and induced genes in L4 neurons at LDR4h. (L) Gene ontology (GO) analysis of genes induced in L2/3 neurons between LDR and LDR30m. (M) GO analysis of genes repressed in L2/3 neurons between LDR30m and LDR2h. (N) GO analysis of genes induced in L4 neurons between LDR and LDR30m. (O) GO analysis of genes repressed in L4 neurons between LDR30m and LDR2h.

To more fully understand the dynamics underlying sensory-dependent transcription, we next asked whether the genes that are induced at LDR30m exhibit sustained expression across the timecourse or whether their expression returned to normal by LDR6h. To do so, we compared the genes that were identified by UniTVelo as induced between LDR and LDR30m to the genes that were repressed between LDR30m and LDR2h for each cell type. Among the 204 genes that were induced in L2/3 neurons between LDR and LDR30m, 112 (55%) were repressed between LDR30m and LDR2h (Fig. 5D). A similar comparison in L4 neurons revealed that 38% of genes induced between LDR and LDR30m are repressed between LDR30m and LDR2h (Fig. 5E). We next compared the dynamics of genes that were upregulated at the LDR4h timepoint, which our data suggests is the peak of LRG programs in both L2/3 and L4 neurons. We observed that, among the 132 genes induced between LDR2h and LDR4h in L2/3 neurons, 98 genes (74%) were repressed between LDR4h and LDR6h (Fig. 5F). The same analysis in L4 neurons revealed that, of the 45 genes that were induced between LDR2h and LDR4h, 30 (67%) were repressed between LDR4h and LDR6h (Fig. 5G). These data suggest that a significant proportion (between 38%-55%) of genes induced at LDR30m are repressed relatively quickly within two hours of induction, while an even more substantial proportion of the genes induced between LDR2h and LDR4h were repressed at LDR6h. These data highlight distinct cohorts of genes in L2/3 and L4 neurons that exhibit transcriptional induction/repression dynamics within the time window captured in our paradigm (Table 3).

**Table 3.** Induced and repressed genes based upon RNA velocity. includes genes identified as transcriptionally induced or repressed in L2/3 and L4 excitatory neurons based upon RNA velocity.

Given that the standard approach for defining sensory-induced genes is to compare RNA abundance between a stimulated and an unstimulated condition (as illustrated in Figs. 3 and 4), we next sought to determine what percentage of the genes that were upregulated at LDR30m based upon total RNA abundance overlapped with the genes identified as induced based upon RNA velocity. As expected, a greater number of genes exhibited heightened expression at LDR30m versus LDR as determined by DEG analysis in both L2/3 and L4 neurons than those identified by velocity as induced in either cell type, suggesting that the list of DEGs for a given cell type in our dataset includes genes that are upregulated via transcription-independent mechanisms, for example possibly due to a decrease in the rate of that gene’s mRNA degradation. However, another interpretation could be that the DEG analysis is more sensitive than the RNA velocity approach. Nevertheless, we found that around 33% of the genes that were upregulated between LDR and LDR30m in L2/3 and L4 neurons based upon DEG analysis were also induced between these time points when assessed by RNA velocity (Fig. 5H,I). A similar analysis of transcripts upregulated between LDR2h and LDR4h revealed 27% overlap for L2/3 but only 11% for L4 neurons, suggesting that LRG programs in L4 neurons might be sustained longer than those in L2/3 neurons (Fig. 5J,K). Consistent with overlap between the results of the DEG and velocity analyses, GO analysis revealed that many of the same or similar ontological categories related to gene function (e.g. *nuclear receptor binding/activity* and *phosphatase binding/activity*) were found among the sets of genes upregulated in L2/3 neurons at LDR30m, with both categories also being among the most strongly enriched genes that are repressed between LDR30m and LDR2h (Fig. 5L,M). A similar pattern emerged for L4 neurons, with GO categories related to *GTPase binding* being induced then repressed (Fig. 4N,O). Thus, the RNA velocity analysis uncovered subsets of DEGs that are most likely to represent *bona fide* IEGs and LRGs based upon their transcriptional dynamics, exhibiting the versatility of the snRNAseq dataset for understanding how sensory experience modifies not just gene expression but transcription explicitly.

### Inference of cell-cell interactions using CellChat uncovers Neurexin and Neuregulin signaling in developing V1

Cells in the brain interact dynamically with one another not only through contact-mediated mechanisms but also through molecular signaling between compatible ligand-receptor pairs. However, a systematic catalog of intercellular interactions in developing visual cortex was lacking. Thus, we next harnessed our snRNAseq dataset to analyze putative cell-cell interactions in V1 across all cell types using the computational tool *CellChat*, which harnesses databases of known ligand-receptor binding partners to estimate the number and strength of putative intercellular communication pathways based upon gene expression data^46^. Given our growing appreciation for the diversity of brain cell types and their functions in circuit development, we applied *CellChat* to map interactions between all cells in our dataset, focusing on the NR condition in which mice are reared normally then processed at the peak of sensory-dependent refinement at P27.

Applying *CellChat* to the NR condition within the snRNAseq dataset, we detected 442 significant ligand-receptor pairs among the 16 cell clusters captured (Fig. 6A,B). We further categorized these pairs as belonging to 58 discrete signaling pathways. Consistent with sensory experience promoting synaptic remodeling and maturation during the time window analyzed, modules related to synapse development and plasticity were among the strongest pathways identified. For example, the Neurexin (Nrxn) family of autism-linked presynaptic adhesion molecules that mediate synapse maintenance and function by binding Neuroligins (Nlgns) and Leucine-rich repeat transmembrane neuronal proteins (Lrrtms) at postsynaptic specializations was the strongest signaling pathway uncovered by *CellChat*. Signaling between Neuregulins (Nrgs) and ErbB receptors, which orchestrates the formation of excitatory synapses onto inhibitory neurons^47,48^, was the second most enriched module identified. Apart from Nrxn and Nrg signaling, Ncam and Cadm (i.e. SynCAM1) adhesion molecules were also identified as active signals in V1. Furthermore, consistent with axonal remodeling occurring during sensory-dependent refinement, EphA and EphB ephrin receptors and Semaphorins 3-6 were also predicted to signal actively (Fig. 6C,D). These findings suggest that cells in V1 work together to shape developing circuits in response to sensory experience via molecular signaling pathways that converge upon synapses.

**Figure 6.**
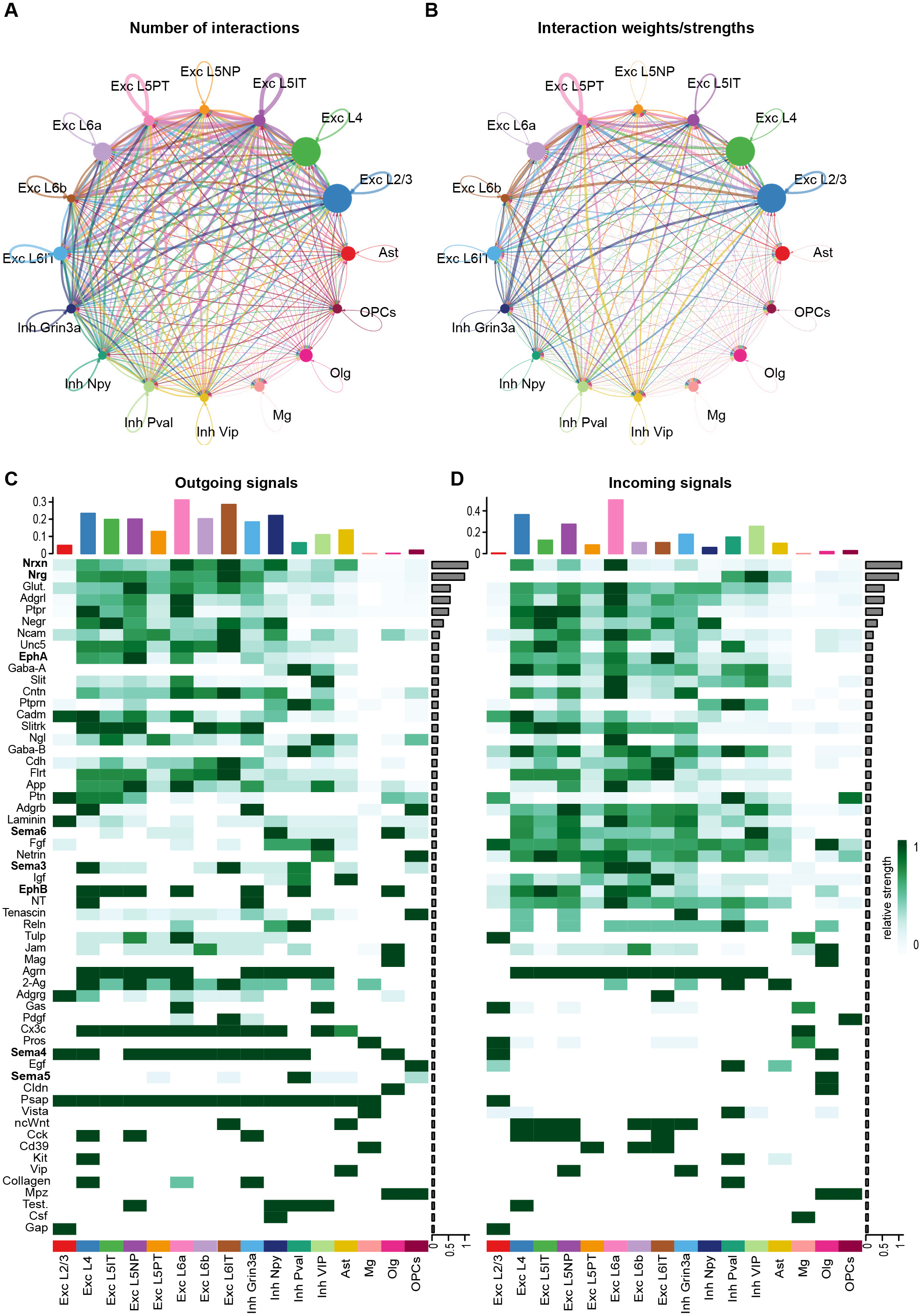
Inference of putative cell:cell interactions in developing V1 using *CellChat*. (A) Cellular communication plot demonstrating the predicted numbers of intercellular ligand-receptor interactions between all cell types in the dataset. (B) Comparative weights/strength of the predicted cell:cell interactions plotted in (A). (C),(D) Heatmaps displaying distinct cell signaling modules (y axis, pathways of interest in bold) predicted by *CellChat* across all cell types (x axis) in the dataset. Top, bars representing the contributions of each cell type to outgoing (C) or incoming (D) signals aggregated across signaling modules. Bar graphs on the right of each heatmap demonstrate the contribution of each individual signaling pathway to the overall interaction score generated in *CellChat*. Heatmap colors indicate the relative strength of a given pathway’s signaling activity as predicted by *CellChat* according to the scale on the right.

We next assessed the putative contributions of the different cell types in V1 to the Nrxn and Nrg signaling pathways identified via *CellChat*. The primary outgoing signals of the Nrxn pathway were Nrxns 3 and 1, and they were most prominently expressed by L6b excitatory neurons (Fig. 6C and 7A). The primary receivers of these signals were Nlgn1 and Lrrtm4, which were most highly expressed in L5-PT neurons but also appeared in L4 neurons and to a lesser extent in other populations as well (Fig. 6D and 7A). In general, we found that excitatory neurons were more heavily involved in both the propagation of outgoing and the receipt of incoming molecular signals than interneurons or glia, with neurons in L6 being particularly active in this regard (Fig. 7A,B). Interestingly, while the inducible gene programs in L2/3 and L4 excitatory neurons shared many features (Fig. 4), these cell classes differed substantially in their participation in cell:cell signaling, with L4 neurons being much more likely to participate in signaling with other V1 cells than L2/3 neurons. Among inhibitory populations, Npy-expressing cells were the strongest senders of outgoing signals while VIP neurons were the strongest receivers (Fig. 7A,B). In contrast, several excitatory populations were predicted to produce Nrg with L6b neurons being the most prominent expressers followed by L4 neurons. All inhibitory cells were relatively strong receivers of Nrg signaling except for VIP neurons (Fig. 7B; Table 4).

**Figure 7.**
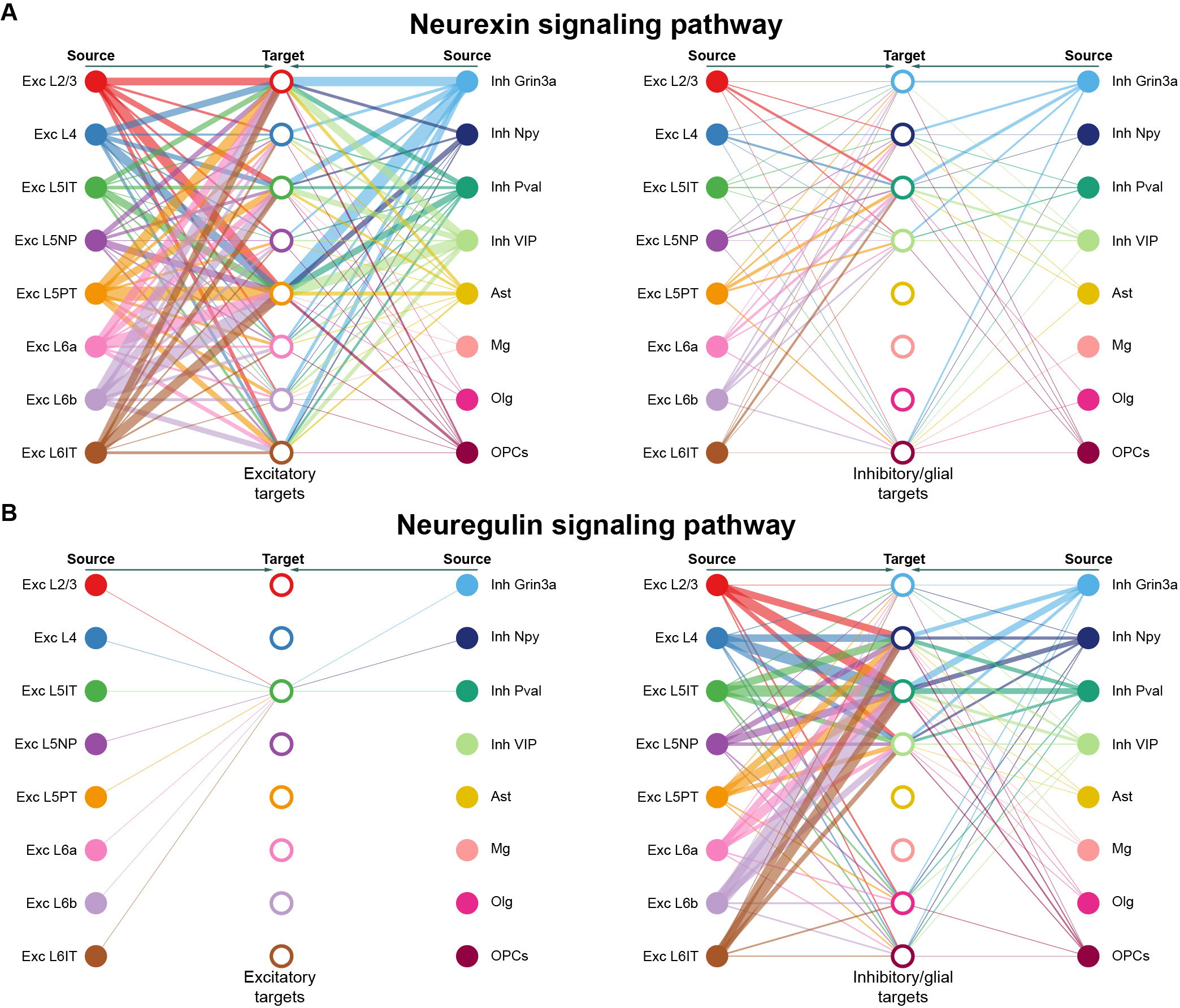
Excitatory-excitatory signaling and excitatory-inhibitory signaling mediated by neurexin and neuregulin pathways, respectively. (A) Hierarchical plot showing Nrxn-mediated interactions from excitatory to excitatory neurons (left) and from excitatory to inhibitory and glial cells (right). (B) Hierarchical plot showing Nrg-mediated interactions from excitatory to excitatory neurons (left) and from excitatory to inhibitory and glial cells (right).

**Table 4.** Cell signaling modules in developing V1 identified by *CellChat*. includes a list of ligand-receptor pairs identified via *CellChat*.

Overall, these data highlight the utility of the snRNAseq resource described here to uncover important principles underlying the molecular control of circuit maturation in the developing brain.

## Discussion

Since the seminal work of Nobel laureates David Hubel and Torsten Wiesel in the 1960s^49,50^, sensory experience has been known to be a major driver of brain development. However, our understanding of the molecular mechanisms engaged by experience to shape brain wiring has remained limited. While molecular adaptations at individual synapses, such as changes in neurotransmitter receptor composition, are well poised to mediate the effects of activity on a neuron’s synapses within an acute time frame, in a developmental context, more global adaptations are warranted. To this point, the idea that robust changes in gene expression driven by sensory stimulation during brain development may play a vital role in circuit refinement is consistent with emerging evidence that neurons in visual cortex undergo significant epigenetic and genomic changes across the first month of life in mice, including between P20 and P27^51,52^. Because these changes in gene expression occur at the cellular rather than the synaptic level, they are likely to exert substantial influence over the development and maintenance of circuits in the long-term.

Inspired by this idea, we here present a whole-transcriptome atlas of sensory-dependent gene expression across 118,529 nuclei representing 16 distinct cell types in the brain. We envision several ways in which this dataset can be used to increase our understanding of brain development. For example, investigators interested in understanding the role(s) of one or more specific genes in brain development can determine whether their genes of interest are expressed in a sensory-dependent manner, and, if so, which cell types upregulate their expression in response to experience. Second, investigators can determine how specific cell types of interest modify their transcriptional profiles in response to sensory stimulation. Finally, given that the dataset includes data from control mice reared normally until P27, investigators can use this data to explore gene expression in V1 in the absence of manipulations of experience.

Several observations that we have made in interrogating this dataset may be of particular interest for future studies. For example, the observation that L2/3 and L4 neurons strongly upregulate intracellular signaling molecules such as protein serine/threonine kinases (including ERK and Sik family members) as early as 30 minutes after stimulation suggests that sensory-dependent gene programs in these cells may reinforce synapse-to-nucleus crosstalk, strengthening the ability of synaptic innervation to shape the neuronal transcriptome. In addition, the observation that excitatory neurons are likely more sensitive to experience than inhibitory cells, both at the level of sensory-induced gene expression changes as well as cell signaling interactions, could increase our understanding of the differential roles that these cell types play in visual function. Among excitatory neurons, the discovery that L2/3 neurons are particularly strongly affected is consistent with a recent study highlighting that the maturation of these cells is influenced visual experience^32^. At the level of cell signaling, our data showing that the strongest signatures were related to Nrxn and Nrg signaling pathways suggests that cellular interactions within developing V1 converge upon synapses. Altogether, we expect this dataset and our experimental investigation thereof to serve as tools for investigators interested in uncovering molecular mechanisms guiding sensory-dependent refinement in the developing brain.

## Methods

### Animals

All experiments were performed in compliance with protocols approved by the Institutional Animal Care and Use Committee at Cold Spring Harbor Laboratory (CSHL). Male C57Bl/6J mice were obtained from the Jackson Laboratory (Cat #000664) then housed at CSHL in an animal facility where average temperatures and humidity were maintained between 68-70° Fahrenheit and 54-58%, respectively. Mice in this study were aged between P18 – P27. Animals had access to food and water *ad libitum*.

### Late-dark-rearing (LDR) paradigm

Male C57Bl/6J mice were obtained from the Jackson laboratory at P18 and allowed to acclimate to the standard 12-hour light/12-hour dark environment of the CSHL animal facility until P20, at which point they were separated into six cohorts. One cohort was maintained under normal housing conditions (normally reared, NR) while the other cohorts were placed inside a well-ventilated,100% light-proof chamber (Actimetrics). Mice in the chamber were housed in complete darkness until P27, at which point one cohort was sacrificed and perfused with ice cold 1X PBS (snRNAseq experiments) or 1X PBS followed by 4% PFA (smFISH experiments) in the dark. The remaining four cohorts of mice were also dark-reared between P20 and P27 but were then re-exposed to light for varying lengths of time: 30 minutes (LDR30m), two hours (LDR2h), four hours (LDR4h), and six hours (LDR6h). After perfusing the mice and removing their brains in the dark, V1 regions were micro-dissected from all cohorts in the wet lab.

### Single-nucleus RNA sequencing and data analysis

#### V1 tissue collection

Whole brains were placed into ice-cold 1X Hank’s Balanced Salt Solution (HBSS) supplemented with Mg^2+^ and Ca^2+^. The V1 brain regions were then bilaterally micro-dissected under a 3.5X-90X Stereo Zoom microscope (AmScope) using a needle blade. Micro-dissected tissue was either immediately processed for snRNAseq or was frozen for later processing.

#### Nuclear suspension preparation

The V1 tissue was transferred to a 1 mL dounce homogenizer containing 300 µL of ice-cold supplemented Homogenization Buffer (0.25M Sucrose, 25mM KCl, 5mM MgCl_2_, 20mM Tricine-KOH, 5mM DTT, 0.75mM Spermine, 2.5mM Spermidine, 0.05X Protease Inhibitor Cocktail, 1U/µL of Rnase Inhibitor and 0.15% IGEPAL CA-630). Note the inclusion of drugs to block gene transcription and protease activity, as well as an RNase inhibitory to protect the integrity of the RNA. The tissue was homogenized with a loose and tight pestle about 10-15 times, respectively. The samples were then filtered using a 20 µm filter.

#### Library construction and sequencing

Single-cell gene expression libraries were prepared using the Single Cell 3′ Gene Expression kit v3.1 (10× Genomics, #1000268) according to manufacturer’s instructions. Libraries were sequenced on an Illumina Nextseq2000 to a mean depth of ∼30,000 reads per cell.

#### Raw data processing

The raw FASTQ files were processed using Cell Ranger (v7.1.0) and aligned to the mm10 reference mouse genome. Loom files for cell dynamics analysis were generated using Velocyto (v0.17.17) by mapping BAM files to the gene annotation GTF file (refdata-gex-mm10-2020-A). Each library derived from the single-nucleus datasets underwent identical processing, resulting in a gene expression matrix of mRNA counts across genes and individual nuclei. Each cell was annotated with the sample name for subsequent batch correction and meta-analysis.

#### Quality control, cell clustering, and cell type annotation

To ensure the integrity of our single-cell RNA sequencing data, we implemented several quality control measures. First, we calculated the log10 of the number of genes per UMI (log10GenesPerUMI), and cells with a value less than 0.85 were excluded. We also removed cells with more than 1% mitochondrial gene expression to reduce noise from apoptotic or damaged cells. Additional thresholds included excluding cells with fewer than 500 UMIs or 300 genes to eliminate low-quality or empty droplets. Doublets were identified and excluded using the DoubletFinder package, with optimal pK values determined for each sample through a sweep analysis^25^. Ambient RNA was removed with DecontX^27^. Following these steps, we applied the standard Seurat (v4) pipeline for data pre-processing (https://satijalab.org/seurat/articles/get_started.html), which included selecting the top 3,000 highly variable genes and regressing out UMI counts and mitochondrial gene percentage for cell clustering.

Clustering utilized the functions *FindNeighbors* and *FindClusters* from Seurat, employing resolutions ranging from 0.1 to 0.5. A resolution of 0.5 was ultimately selected for clustering. To identify major cell types, the *ConserveredMarkers* function (log2 fold change > 0.25, MAST test, adjusted p-value < 0.05 with Bonferroni correction), with pct.1 > 70% and pct.2 < 30% identified unique and highly enriched differentially expressed genes (DEGs) in specific clusters compared to others. Cell types were manually annotated based on the expression of conserved markers^21,28^, ensuring precise identification and accurate analysis of cellular phenotypes.

#### DEG analysis

Differentially expressed genes (DEGs) between conditions were identified using the DEseq2 function within Seurat v4. DEGs were identified for each of the 16 individual clusters included in the dataset.

#### RNA velocity analysis

Cell velocity analysis was conducted on L2/3 and L4 excitatory neurons using the UniTVelo (v0.2.4) tool within the scVelo (v0.2.5.) framework, focusing on the 2000 most variably expressed genes. Genes were categorized based on their fit_t scores such that those with a fit_t>0 were classified as induced genes, whereas genes with a fit_t <0 were identified as repressed genes.

#### Cell-cell interaction analysis

The R package *CellChat* (http://www.cellchat.org/) was utilized to infer cell-cell interactions within our dataset. We adhered to the standard pipeline and default parameters set by *CellChat*. The complete CellChatDB.mouse database was employed, which categorizes ligand-receptor pairs into “Secreted Signaling,” “ECM-Receptor,” and “Cell-Cell Contact.” Additionally, we conducted CellChat analyses on the overall dataset and separately for conditions at specific timepoints—LDR0, LDR30m, LDR2h, LDR4h, LDR6h, and NR, although we focus on the NR condition in this paper.

#### Enrichment analysis

Gene Ontology (GO) enrichment analysis was conducted using the “clusterProfiler”(v4.10.0) package. For the analysis of differentially expressed genes (DEGs), only genes with an adjusted p-value less than 0.05 and a log2 fold change greater than log2(1.5) were included. For the analysis of induced and repressed genes, all identified genes were considered. The parameters for the GO analysis were set with a p-value cutoff of 0.05 and a q-value cutoff of 0.2, using the Benjamini-Hochberg (BH) method for adjusting p-values. This approach ensures rigorous identification of biological processes significantly associated with the gene sets under study.

#### Real-time qPCR

Flash frozen V1 samples were processed for RNA extraction using Trizol (ThermoFisher cat #15596018) according to the manufacturer’s protocol. The cDNA library was built using iScript Kit (BioRad, cat #1725037) and Oligo d(T) primers. The real-time PCR were performed using SybrGreen kit (Fisher, cat #A25742) and standard PCR temperature protocol. *Fos* and *Jun* expression were normalized to *Gapdh* levels. The following primer sequences were used.

*Fos* (Forward): 5’-GGGAATGGTGAAGACCGTGTCA-3’

*Fos* (Reverse): 5’-GCAGCCATCTTATTCCGTTCCC-3’

*Jun* (Forward): 5’-CAGTCCAGCAATGGGCACATCA-3’

*Jun* (Reverse): 5’-GGAAGCGTGTTCTGGCTATGCA-3’

*Gapdh* (Forward): 5’-CATCACTGCCACCCAGAAGACTG-3’

*Gapdh* (Reverse): 5’-ATGCCAGTGAGCTTCCCGTTCAG-3’

#### Single-molecule fluorescence in situ hybridization (smFISH)

Animals were anesthetized with a ketamine and xylazine cocktail (K: 90 mg/kg X: 10 mg/kg) before perfusion with ice-cold phosphate-buffered saline (PBS) followed by 4% paraformaldehyde (PFA) in 1X PBS. Brains were then drop-fixed in 4% PFA in 1X PBS for 24 hours. Brains were then washed with 1X PBS thrice for 10 min before being transferred to a 30% sucrose solution at 4° C. After dehydration, brains were embedded in optimal cutting temperature (OCT; VWR cat #25608-930) and stored at −80° C. 20-μm thick coronal sections containing the visual cortex were cut using a cryostat and thaw-mounted onto a Superfrost Plus microscope slide (Thermo Fisher Scientific, cat #1255015) and stored at −80° C until the experiment. FISH was performed using the RNAScope platform V2 kit (Advanced Cell Diagnostics (ACD), cat #323100) according to the manufacturer’s protocol for fixed-frozen sections. Samples were then counterstained with DAPI before ProLong Gold Antifade was applied. A 1.5X thickness coverslip was then applied to the slides which were then stored at 4° C until imaging. Commercial probes from ACDBio were obtained to detect the following genes: *Fos* (316921), *Nr4a1* (423342-C2)*, and Jun* (453561-C3).

#### Confocal Imaging

smFISH images were acquired using the Zeiss LSM780 with a x20/0.8 objective. Z-stack images were acquired.

#### FISH Quantification

FISH images were analyzed using FIJI. For each image, ROIs of layer 4 and layer 2/3 of the visual cortex were defined. The mean gray values were then taken for each ROI. For each mouse, the average mean gray value across both hemispheres was analyzed for both layer 4 and layer 2/3. A 2-way ANOVA was performed to test for significance.

## Data availability

Both raw and processed snRNA-seq data are available at Gene Expression Omnibus under accession number GSE269482.

## Acknowledgements

We thank Dr. Timothy J. Burbridge (Boston Children’s Hospital), Dr. Gabrielle Pouchelon (Cold Spring Harbor Laboratory), Dr. Leena Al Ibrahim (King Abdulla University of Science and Technology), Dr. Marty Yang (University of California, San Francisco), and members of the Cheadle lab for critical input and feedback on the manuscript. This work was supported by the following grants (to L.C.): R00MH120051, DP2MH132943, R01NS131486, Rita Allen Scholar Award, McKnight Scholar Award, Klingenstein-Simons Fellowship Award in Neuroscience, and a Brain and Behavior Foundation NARSAD grant. Dr. Cheadle is a Howard Hughes Medical Institute Freeman Hrabowski Scholar.

## Author contribution statement

AMX and LC conceptualized the study. AMX and CK designed, optimized, and performed snRNAseq. AMX performed and analyzed qPCR. CK performed and analyzed fluorescence *in situ* hybridization. QL and AMX analyzed the snRNAseq data, and QL and LC generated figures. LC wrote the paper.

